## Supplementary material for "Epigenetic drift during long-term culture of cells *in vitro*": Combined PDF with all supplemental figures (S1-S5) and supplemental tables S1, S3, S4, S5, S6, S7, S8, S9, and S10.

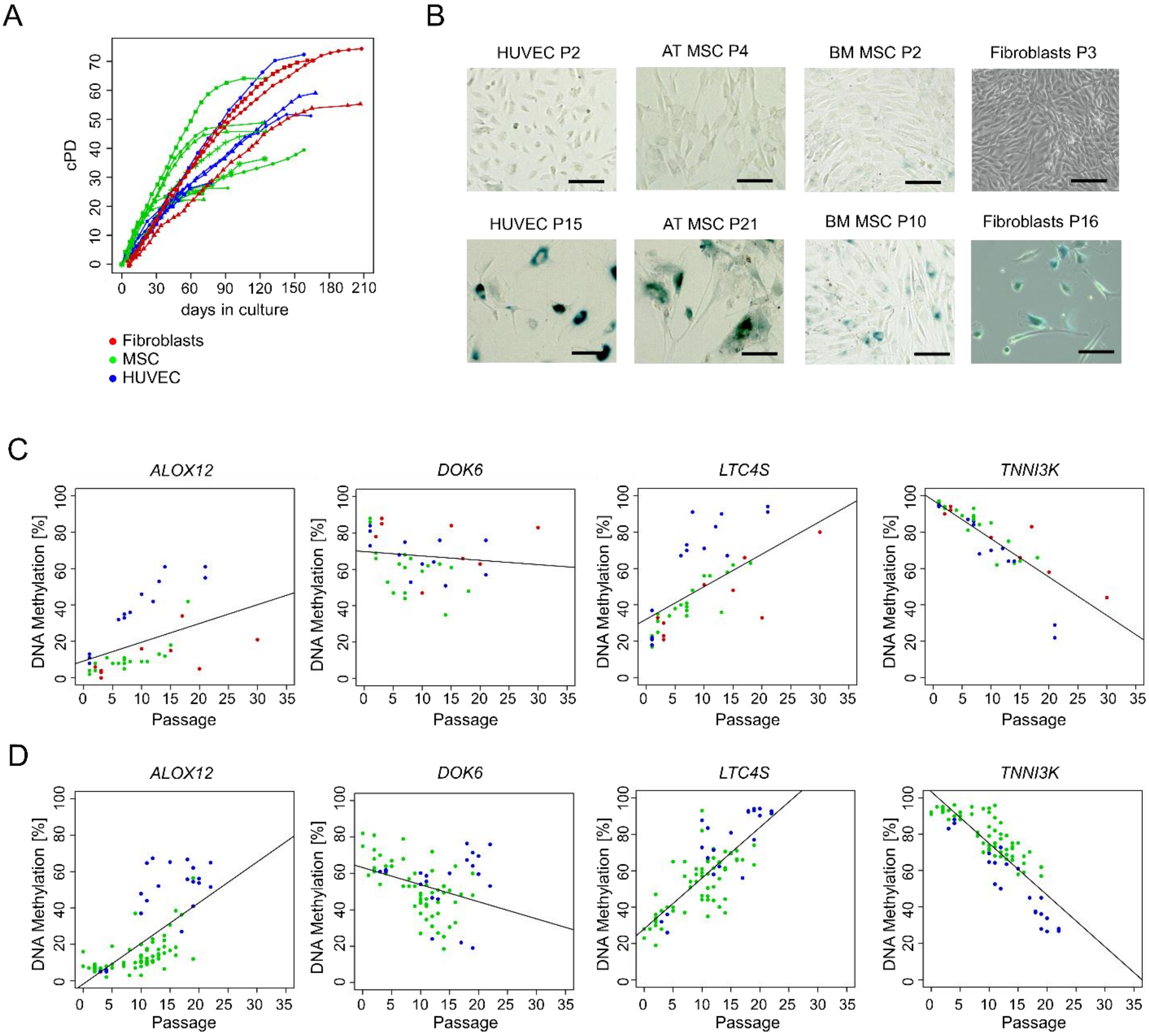


**Supplemental Figure S1: Growth curves and long-term culture-associated changes in cell preparations. A)** Growths curves of the cell preparations of the training set (each dot represents a passage; cPD = cumulative population doublings). **B)** Representative images of senescence-associated beta galactosidase staining (AT = adipose-tissue-derived, BM = bone-marrow-derived, P = passage, size bar = 100 µm). DNAm levels were measured by pyrosequencing in the training **(C)**, and validation set **(D).** DNAm at all of the four long-term culture-associated CpGs correlated with passage numbers of cell preparations.

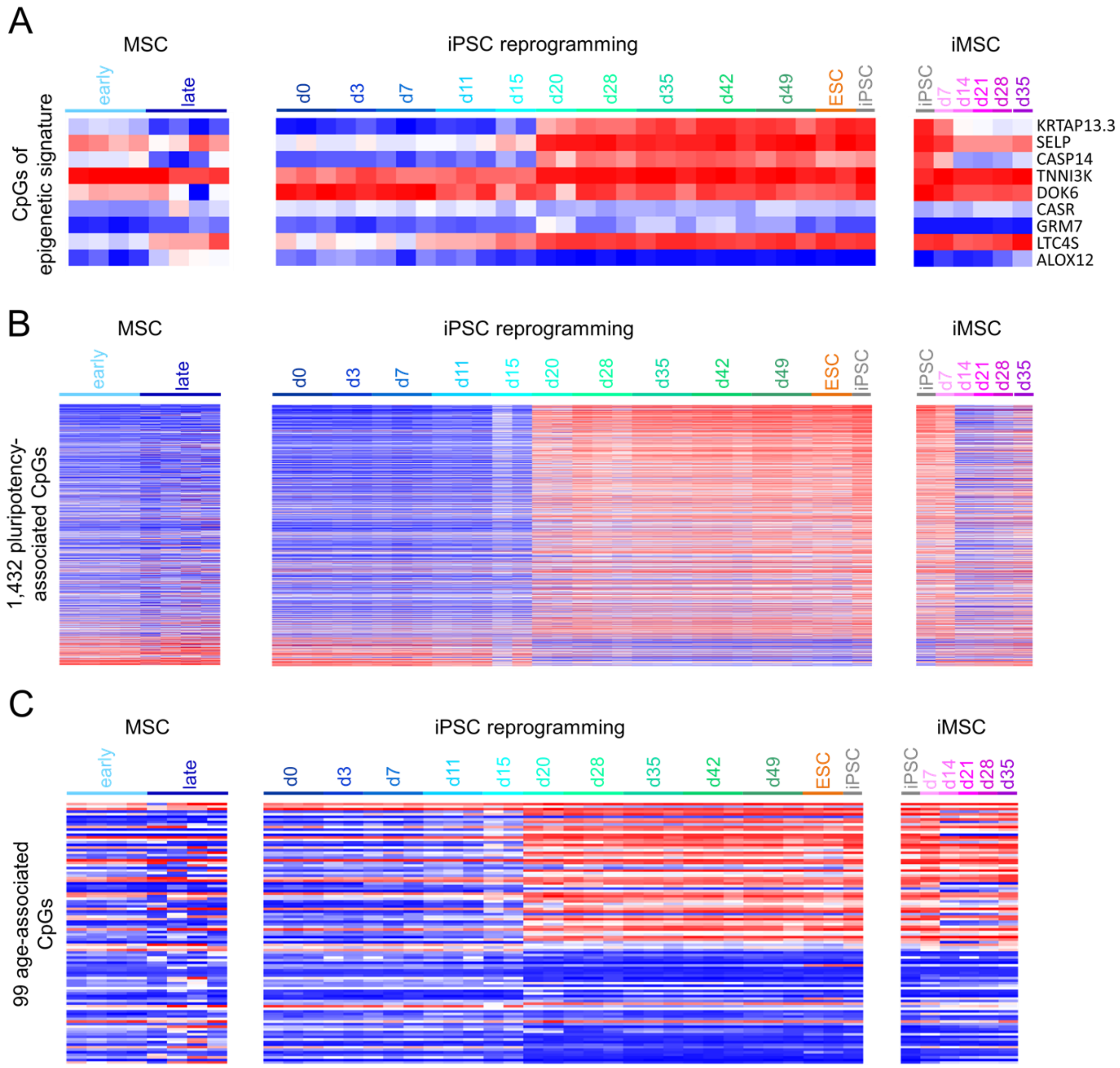


**Supplemental Figure S2: DNAm changes of different CpG subsets are reset at day 20 during reprogramming into iPSCs.** DNAm levels at **A)** 9 CpGs of our epigenetic signatures (1), **B)** 1,432 pluripotency associated CpG sites (2), and **C)** 99 age-associated CpGs of our previously described age-predictor for blood samples (3) are depicted in MSCs of early (passage 2) and late passages (passage 7 to 16; GSE37067, left panel) (4), during reprogramming of fibroblasts into iPSCs (GSE54848; central panel), (5) and re-differentiation of iPSCs to iMSCs (GSE54767; right panel) (6).

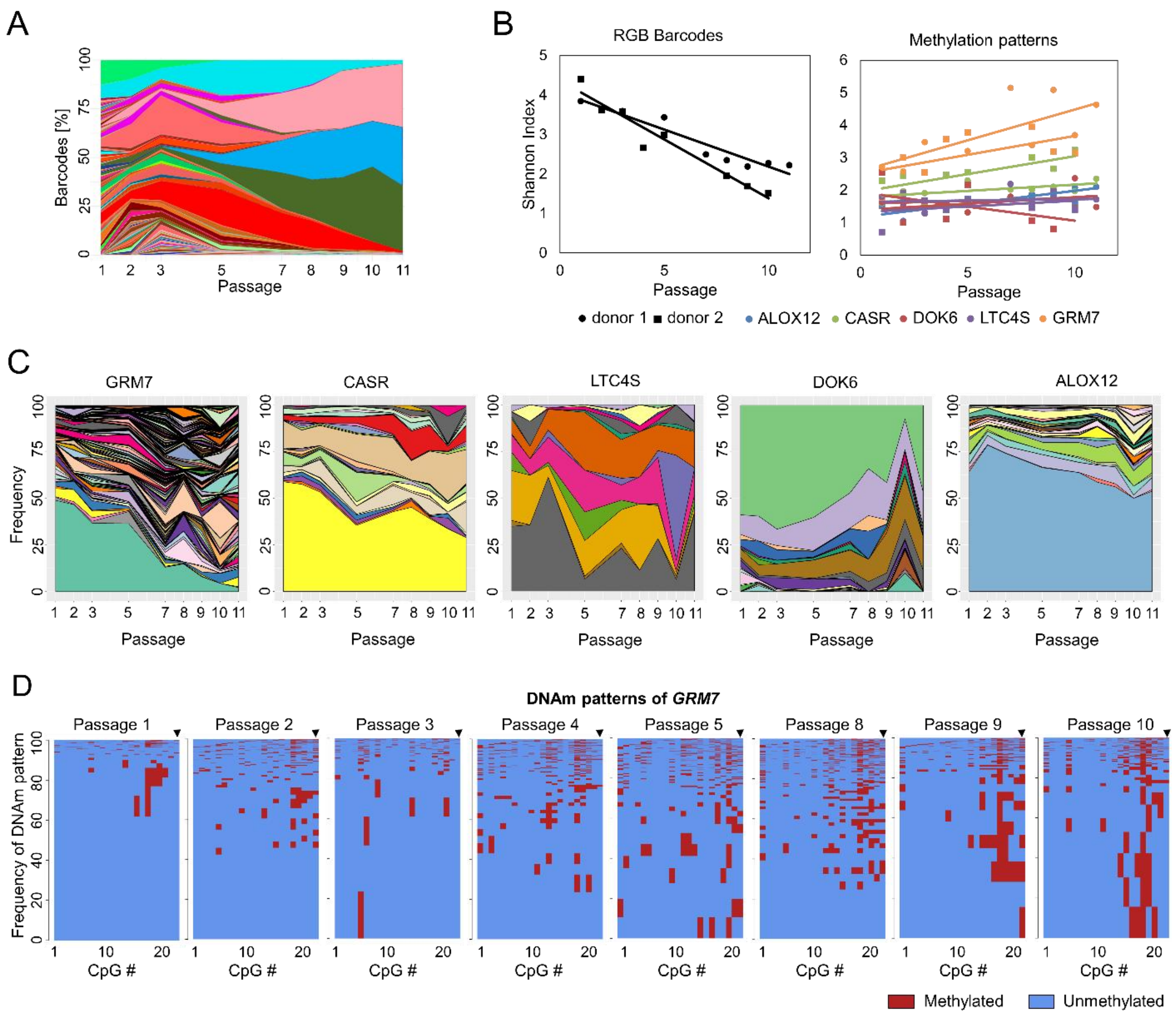


**Supplemental Figure S3: Variations of DNAm patterns and estimation of Shannon index. A)** Deep sequencing analysis of random barcodes demonstrates that MSCs of the second preparation become oligoclonal at later passages (in analogy to Figure 3A). **B)** Shannon index of RGB barcodes in the two UC-MSC donors and of the DNAm patterns in five different amplicons of the two donors. **C)** Frequencies of different DNAm patterns within neighboring CpGs for the second MSC preparation (in analogy to Figure 3C). **D)** Individual DNAm patterns are exemplarily depicted for 22 neighboring CpGs of *GRM7* of UC-MSC donor 1 (corresponding to Figure 3). The frequency of DNAm patterns resembles the percentage of corresponding reads in deep sequencing analysis. The height of each pattern corresponds to the frequency of this specific DNAm pattern, while the sum of all pattern frequencies adds up to 100%.

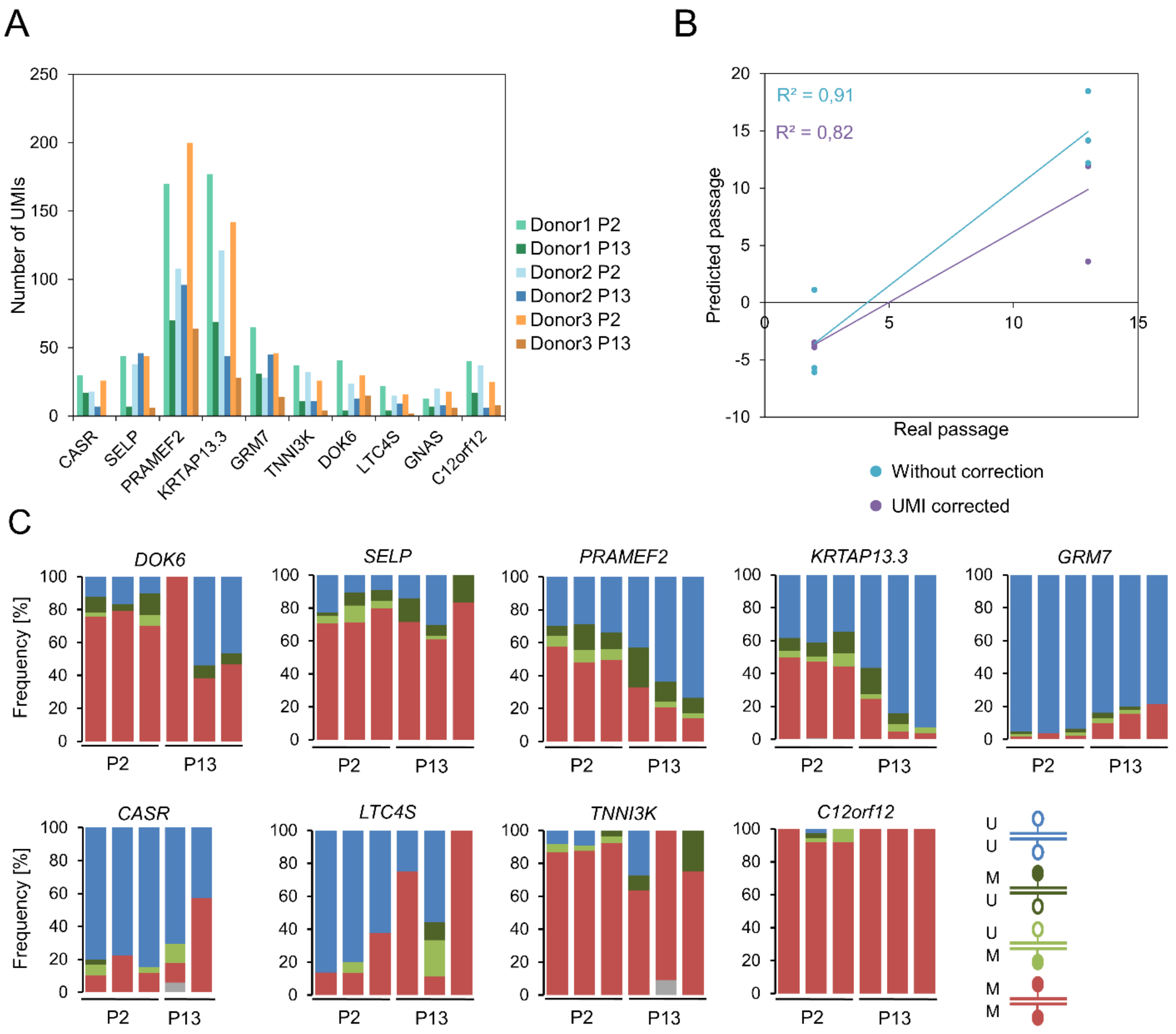


**Supplemental Figure S4: Analysis of hairpin BBA-Seq. A)** Number of different unique molecular identifiers (UMIs) within hairpin loops detected for each amplicon. **B)** Passage predictions based on eight culture-associated CpGs either without or with consideration of different UMIs. **C)** Frequency of homo- and hemimethylation at culture-expansion-associated CpG sites that were used for predictions of passage numbers and at a highly methylated region (*C12orf12*). Grey colored regions resemble sequencing errors at one of the complementary CpG sites. One late passage donor of *CASR* is missing due to failed sequencing.

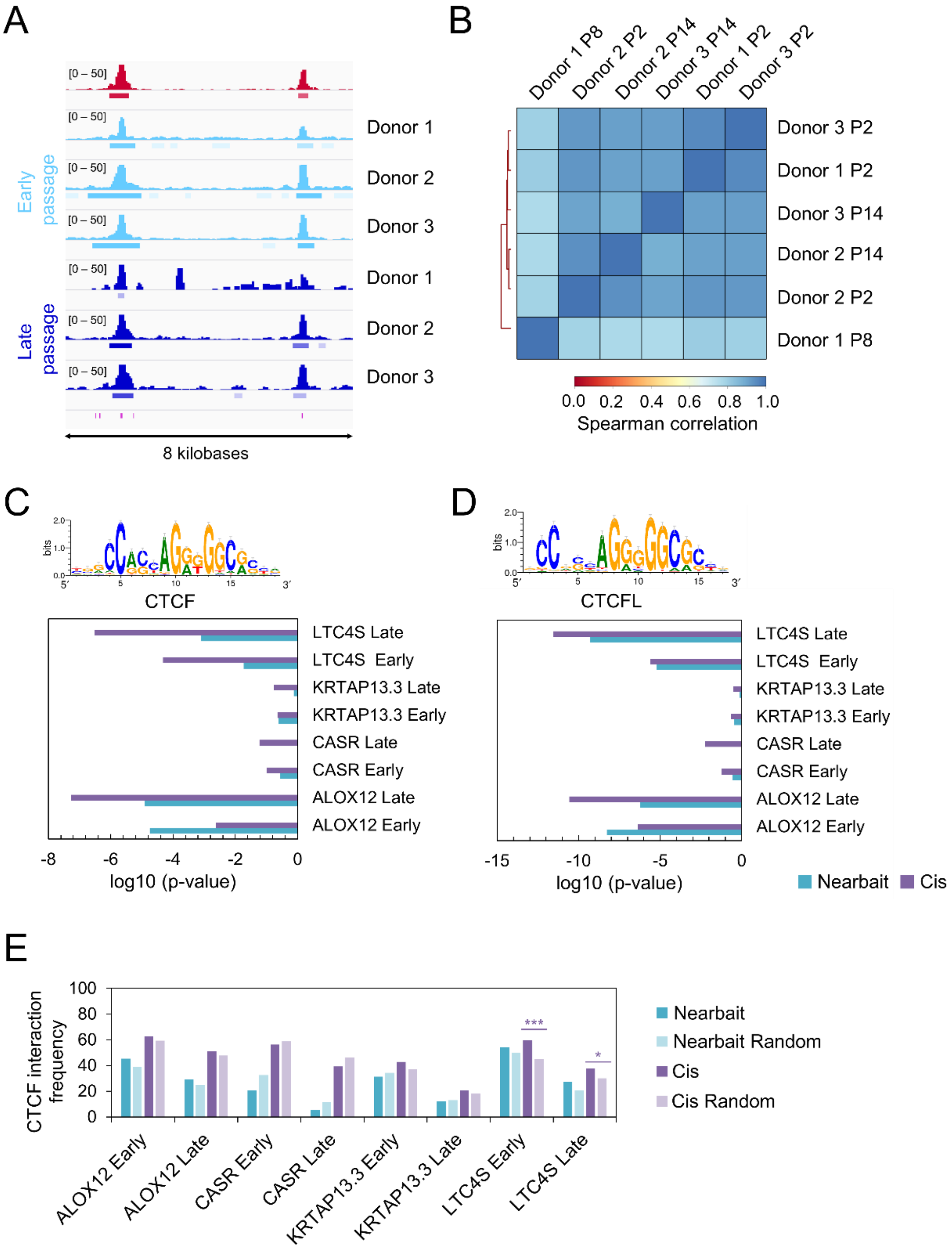


**Supplemental Figure S5: Analysis of CTCF binding in early and late passage cells. A)** Representative Integrative Genomics Viewer (IGV) overview of ChIP-seq peaks in early (P2, light blue) and late (P8 – P14, dark blue) passages. Peaks are associated to predicted CTCF motifs (violet lines at the bottom) and are reproducible in several donors, both in early and late passages as well as in a publicly available dataset (red, embryonic stem cell derived MSCs) (7). **B)** Spearman correlation of normalized CTCF ChIP-seq peaks reveals high correlation for all samples irrespective of their passage number. **C, D)** Enrichment analysis of predicted motif binding sites for CTCF (C) and CTCFL (D) in high interacting regions of the 4C experiment. P-values were calculated with the RGT motif enrichment tool. Significant enrichment was particularly observed for the hypermethylated CpGs in *LTC4S* and *ALOX12*. **E)** Enrichment of CTCF ChIP-seq peaks within high interacting sites of the four baits in the 4C experiment compared to random background regions. To this end, CTCF binding peaks were further divided into “early” or “late” CTCF peaks, derived from corresponding early or late passaged samples respectively. Significance was tested by Fisher’s exact test (* $p < 0.05$; ** $p < 0.01$; *** $p < 0.001$).

**Supplemental Table S1. Illumina 450k BeadChip datasets.**

| Geo accession | GSE number | cell type | passage | Publication |
|---|---|---|---|---|
| GSM1004625 | GSE40927 | fibroblast | 4 | Kurian, Sancho-Martinez (8) |
| GSM1004626 | GSE40927 | fibroblast | 5 | Kurian, Sancho-Martinez (8) |
| GSM1004627 | GSE40927 | fibroblast | 14 | Kurian, Sancho-Martinez (8) |
| GSM1004643 | GSE40927 | HUVEC | 4 | Kurian, Sancho-Martinez (8) |
| GSM1004644 | GSE40927 | HUVEC | 4 | Kurian, Sancho-Martinez (8) |
| GSM1004645 | GSE40927 | HUVEC | 17 | Kurian, Sancho-Martinez (8) |
| GSM1027664 | GSE41933 | MSC | 2 | Reinisch, Etchart (9) |
| GSM1027665 | GSE41933 | MSC | 2 | Reinisch, Etchart (9) |
| GSM1027666 | GSE41933 | MSC | 2 | Reinisch, Etchart (9) |
| GSM1027667 | GSE41933 | MSC | 2 | Reinisch, Etchart (9) |
| GSM1027668 | GSE41933 | MSC | 2 | Reinisch, Etchart (9) |
| GSM1027669 | GSE41933 | MSC | 2 | Reinisch, Etchart (9) |
| GSM1027670 | GSE41933 | MSC | 2 | Reinisch, Etchart (9) |
| GSM1027671 | GSE41933 | MSC | 2 | Reinisch, Etchart (9) |
| GSM1027672 | GSE41933 | MSC | 2 | Reinisch, Etchart (9) |
| GSM1027673 | GSE41933 | fibroblast | 2 | Reinisch, Etchart (9) |
| GSM1027674 | GSE41933 | fibroblast | 2 | Reinisch, Etchart (9) |
| GSM1027675 | GSE41933 | fibroblast | 2 | Reinisch, Etchart (9) |
| GSM853409 | GSE37066 | MSC | 2 | Koch, Reck (4) |
| GSM853410 | GSE37066 | MSC | 2 | Koch, Reck (4) |
| GSM853411 | GSE37066 | MSC | 2 | Koch, Reck (4) |
| GSM853412 | GSE37066 | MSC | 2 | Koch, Reck (4) |
| GSM853413 | GSE37066 | MSC | 3 | Koch, Reck (4) |
| GSM909610 | GSE37066 | MSC | 7 | Koch, Reck (4) |
| GSM909612 | GSE37066 | MSC | 12 | Koch, Reck (4) |
| GSM909614 | GSE37066 | MSC | 16 | Koch, Reck (4) |
| GSM909616 | GSE37066 | MSC | 14 | Koch, Reck (4) |
| GSM909618 | GSE37066 | MSC | 14 | Koch, Reck (4) |
| GSM3230133 | GSE116375 | MSC | 4 | this study |
| GSM3230134 | GSE116375 | MSC | 4 | this study |
| GSM3230135 | GSE116375 | MSC | 4 | this study |
| GSM3230136 | GSE116375 | MSC | 10 | this study |
| GSM3230137 | GSE116375 | MSC | 10 | this study |
| GSM3230138 | GSE116375 | MSC | 10 | this study |
| GSM2340959 | GSE87797 | MSC | 2 | Fernandez-Rebollo, Mentrup (10) |
| GSM2340960 | GSE87797 | MSC | 2 | Fernandez-Rebollo, Mentrup (10) |
| GSM2340961 | GSE87797 | MSC | 2 | Fernandez-Rebollo, Mentrup (10) |
| GSM2340962 | GSE87797 | MSC | 2 | Fernandez-Rebollo, Mentrup (10) |
| GSM2340963 | GSE87797 | MSC | 2 | Fernandez-Rebollo, Mentrup (10) |
| GSM2340964 | GSE87797 | MSC | 2 | Fernandez-Rebollo, Mentrup (10) |
| GSM2340965 | GSE87797 | MSC | 2 | Fernandez-Rebollo, Mentrup (10) |
| GSM2340966 | GSE87797 | MSC | 2 | Fernandez-Rebollo, Mentrup (10) |
| GSM2340967 | GSE87797 | MSC | 2 | Fernandez-Rebollo, Mentrup (10) |
| GSM2340968 | GSE87797 | MSC | 2 | Fernandez-Rebollo, Mentrup (10) |

| | | | | |
|---|---|---|---|---|
| GSM2340969 | GSE87797 | MSC | 2 | Fernandez-Rebollo, Mentrup (10) |
| GSM2340970 | GSE87797 | MSC | 2 | Fernandez-Rebollo, Mentrup (10) |
| GSM1347975 | GSE55888 | MSC | 4 | Schellenberg, Joussen (11) |
| GSM1347978 | GSE55888 | MSC | 4 | Schellenberg, Joussen (11) |
| GSM1347981 | GSE55888 | MSC | 4 | Schellenberg, Joussen (11) |
| GSM1347984 | GSE55888 | MSC | 4 | Schellenberg, Joussen (11) |
| GSM1347987 | GSE55888 | MSC | 4 | Schellenberg, Joussen (11) |
| GSM1347989 | GSE55888 | MSC | 4 | Schellenberg, Joussen (11) |
| GSM1347991 | GSE55888 | MSC | 4 | Schellenberg, Joussen (11) |
| GSM1347993 | GSE55888 | MSC | 4 | Schellenberg, Joussen (11) |
| GSM2186934 | GSE82234 | HUVEC | 4 | Franzen, Zirkel (12) |
| GSM2186935 | GSE82234 | HUVEC | 4 | Franzen, Zirkel (12) |
| GSM2186936 | GSE82234 | HUVEC | 4 | Franzen, Zirkel (12) |
| GSM3230129 | GSE116375 | HUVEC | 10 | this study |
| GSM3230130 | GSE116375 | HUVEC | 15 | this study |
| GSM3230131 | GSE116375 | HUVEC | 12 | this study |
| GSM3230132 | GSE116375 | HUVEC | 8 | this study |
| GSM2186937 | GSE82234 | HUVEC | 20 | Franzen, Zirkel (12) |
| GSM2186938 | GSE82234 | HUVEC | 18 | Franzen, Zirkel (12) |
| GSM2186939 | GSE82234 | HUVEC | 13 | Franzen, Zirkel (12) |

**Supplemental table S2: Long-term culture-associated CpGs with cutoff R > 0.7 or R < -0.7**. This list of CpG sites is provided as additional Excel file.

**Supplemental Table S3. Thirty selected long-term culture-associated CpGs**

| Illumina ID | Pearson (R) | $R^2$ | Slope (m) |
|---|---|---|---|
| cg25281820 | -0.87786992 | 0.770655597 | -0.02079236 |
| cg03421657 | -0.86871368 | 0.754663451 | -0.02217539 |
| cg21567022 | -0.86815882 | 0.753699738 | -0.03263046 |
| cg10588720 | -0.86669937 | 0.751167791 | -0.02130874 |
| cg16470423 | -0.86548811 | 0.749069674 | -0.02626191 |
| cg23054188 | -0.86328129 | 0.74525459 | -0.02776338 |
| cg05264232 | -0.86248453 | 0.743879567 | -0.02051906 |
| cg07403610 | -0.86147656 | 0.742141858 | -0.024732 |
| cg17625256 | -0.8603098 | 0.740132952 | -0.0238157 |
| cg00063346 | -0.85976139 | 0.73918965 | -0.02255066 |
| cg25968937 | -0.85690853 | 0.734292227 | -0.03551924 |
| cg00791548 | -0.85496971 | 0.730973203 | -0.02079242 |
| cg07566463 | -0.85404073 | 0.729385561 | -0.02079289 |
| cg16819051 | -0.85340441 | 0.728299093 | -0.02279132 |
| cg08732456 | -0.85092846 | 0.724079248 | -0.02796559 |
| cg04682775 | 0.80592194 | 0.649510173 | 0.02106277 |
| cg19443920 | 0.80854957 | 0.653752415 | 0.02610184 |
| cg17180284 | 0.80937209 | 0.655083181 | 0.02006929 |
| cg24628744 | 0.80950526 | 0.655298762 | 0.02028689 |
| cg06221449 | 0.81021837 | 0.656453811 | 0.02305057 |
| cg01815912 | 0.81744955 | 0.668223773 | 0.02761235 |
| cg19759135 | 0.81834739 | 0.669692446 | 0.02261348 |
| cg05054998 | 0.81942121 | 0.671451112 | 0.03632707 |
| cg23602058 | 0.82959135 | 0.688221809 | 0.02256118 |
| cg26683398 | 0.84259943 | 0.709973806 | 0.02118855 |
| cg03762994 | 0.84439666 | 0.713005722 | 0.02743842 |
| cg02717339 | 0.84612683 | 0.715930609 | 0.0210322 |
| cg27456203 | 0.85926672 | 0.738339289 | 0.02156726 |
| cg11394785 | 0.86553634 | 0.749153148 | 0.02060183 |
| cg04065086 | 0.8684504 | 0.754206094 | 0.02188371 |

**Supplemental Table S4. Pyrosequencing primers**

| gene name | primer | Sequence 5’ -> 3’ |
|---|---|---|
| *ALOX12* | forward | GGGGTTATTTTTAATTTTTAAAGGAT |
| *ALOX12* | reverse | Biotin - AAAACACAACCAATCCCCACAA |
| *ALOX12* | sequencing | TTATTTTTAATTTTTAAAGGA |
| *DOK6* | forward | Biotin-ATGAGAAAATTGAGATATAATTTTATTTAGGAAATAG |
| *DOK6* | reverse | CACCTTTTCTTCTAAAATCTACAAATCCC |
| *DOK6* | sequencing | TCACACTCAAAATCTAACCTA |
| *LTC4S* | forward | TTTTAGGGTTTTGTAGATTTTTATATTATGTTGGAGTTAG |
| *LTC4S* | reverse | Biotin - CACCCAAAAAACCTTAAACAAATTTCC |
| *LTC4S* | sequencing | AATTTTTGTTAAATTTTTTTTT |
| *TNNI3K* | forward | Biotin- ATTTTGTGGTTTTTTATAATGTTTTAGGAGTGTGATAA |
| *TNNI3K* | reverse | CCATAATCACTTTATTCACTACATCACCAATACCCATTC |
| *TNNI3K* | sequencing | CAATAAAATACCTAACATAATACT |

**Supplemental Table S5. Pyrosequencing results of training dataset**

| Sample | *ALOX12* | *DOK6* | *LTC4S* | *TNNI3K* | Real passage | Predicted passage |
|---|---|---|---|---|---|---|
| AT MSC 1 | 0.04 | 0.88 | 0.23 | 0.95 | 1 | 3 |
| AT MSC 1 | 0.09 | 0.66 | 0.48 | 0.84 | 8 | 8 |
| AT MSC 1 | 0.18 | 0.61 | 0.62 | 0.64 | 15 | 16 |
| AT MSC 2 | 0.04 | 0.86 | 0.23 | 0.97 | 1 | 2 |
| AT MSC 2 | 0.09 | 0.59 | 0.56 | 0.85 | 10 | 8 |
| AT MSC 2 | 0.42 | 0.48 | 0.63 | 0.66 | 18 | 12 |
| AT MSC 3 | 0.02 | 0.88 | 0.17 | 0.96 | 1 | 2 |
| AT MSC 3 | 0.05 | 0.61 | 0.39 | 0.89 | 7 | 6 |
| AT MSC 3 | 0.12 | 0.35 | 0.58 | 0.63 | 14 | 16 |
| Fibroblast 1 | 0.06 | 0.78 | 0.33 | 0.9 | 2 | 5 |
| Fibroblast 1 | 0.16 | 0.47 | 0.51 | 0.77 | 10 | 10 |
| Fibroblast 1 | 0.34 | 0.66 | 0.66 | 0.83 | 17 | 7 |
| Fibroblast 2 | 0.04 | 0.85 | 0.3 | 0.92 | 3 | 4 |
| Fibroblast 2 | 0.2 | 0.85 | 0.59 | 0.32 | 37 | 28 |
| Fibroblast 3 | 0.03 | 0.88 | 0.23 | 0.94 | 3 | 3 |
| Fibroblast 3 | 0.15 | 0.84 | 0.48 | 0.66 | 15 | 14 |
| Fibroblast 3 | 0.21 | 0.83 | 0.8 | 0.44 | 30 | 24 |
| Fibroblast 4 | 0 | 0.85 | 0.21 | 0.92 | 3 | 4 |
| Fibroblast 4 | 0.05 | 0.63 | 0.33 | 0.58 | 20 | 18 |
| HUVEC 1 | 0.13 | 0.73 | 0.37 | 0.94 | 1 | 3 |
| HUVEC 1 | 0.36 | 0.53 | 0.91 | 0.68 | 8 | 14 |
| HUVEC 1 | 0.61 | 0.51 | 0.67 | 0.64 | 14 | 11 |
| HUVEC 2 | 0.13 | 0.81 | 0.22 | 0.94 | 1 | 2 |
| HUVEC 2 | 0.32 | 0.68 | 0.67 | 0.87 | 6 | 5 |
| HUVEC 2 | 0.46 | 0.63 | 0.71 | 0.7 | 10 | 11 |
| HUVEC 3 | 0.08 | 0.81 | 0.21 | 0.95 | 1 | 2 |
| HUVEC 3 | 0.33 | 0.75 | 0.7 | 0.84 | 7 | 7 |
| HUVEC 3 | 0.42 | 0.64 | 0.83 | 0.71 | 12 | 11 |
| HUVEC 3 | 0.55 | 0.57 | 0.94 | 0.29 | 21 | 27 |
| HUVEC 4 | 0.11 | 0.84 | 0.18 | 0.95 | 1 | 2 |
| HUVEC 4 | 0.35 | 0.68 | 0.73 | 0.86 | 7 | 6 |
| HUVEC 4 | 0.53 | 0.76 | 0.9 | 0.64 | 13 | 13 |
| HUVEC 4 | 0.61 | 0.76 | 0.91 | 0.22 | 21 | 29 |
| BM MSC 1 | 0.08 | 0.66 | 0.31 | 0.93 | 2 | 4 |
| BM MSC 1 | 0.11 | 0.47 | 0.41 | 0.93 | 7 | 4 |
| BM MSC 2 | 0.04 | 0.66 | 0.25 | 0.94 | 2 | 3 |
| BM MSC 2 | 0.08 | 0.44 | 0.34 | 0.87 | 7 | 6 |
| BM MSC 2 | 0.13 | 0.63 | 0.36 | 0.75 | 13 | 10 |
| BM MSC 3 | 0.08 | 0.69 | 0.35 | 0.93 | 2 | 4 |
| BM MSC 3 | 0.08 | 0.63 | 0.4 | 0.81 | 6 | 9 |
| BM MSC 3 | 0.09 | 0.62 | 0.56 | 0.62 | 11 | 17 |
| BM MSC 2 | 0.1 | 0.68 | 0.37 | 0.88 | 7 | 6 |
| BM MSC 3 | 0.08 | 0.47 | 0.38 | 0.89 | 5 | 6 |
| BM MSC 1 | 0.11 | 0.53 | 0.34 | 0.92 | 4 | 4 |

Methylation values of single CpGs are depicted as β-values ranging from 0 to 1.

**Supplemental Table S6. Pyrosequencing results of validation dataset**

| Sample | ALOX12 | DOK6 | LTC4S | TNNI3K | Real passage | Predicted passage |
|---|---|---|---|---|---|---|
| BM MSC 4 | 0.08 | 0.62 | 0.33 | 0.94 | 2 | 3 |
| BM MSC 5 | 0.07 | 0.61 | 0.3 | 0.95 | 2 | 3 |
| BM MSC 6 | 0.08 | 0.68 | 0.34 | 0.94 | 2 | 3 |
| BM MSC 7 | 0.07 | 0.59 | 0.28 | 0.95 | 1 | 3 |
| BM MSC 8 | 0.07 | 0.54 | 0.38 | 0.93 | 3 | 4 |
| BM MSC 9 | 0.09 | 0.63 | 0.32 | 0.95 | 2 | 3 |
| BM MSC 10 | 0.07 | 0.63 | 0.36 | 0.92 | 2 | 4 |
| BM MSC 11 | 0.05 | 0.73 | 0.47 | 0.94 | 2 | 4 |
| BM MSC 5 | 0.11 | 0.38 | 0.82 | 0.84 | 10 | 10 |
| BM MSC 6 | 0.03 | 0.49 | 0.59 | 0.71 | 10 | 14 |
| BM MSC 7 | 0.13 | 0.27 | 0.57 | 0.79 | 10 | 10 |
| BM MSC 8 | 0.09 | 0.42 | 0.44 | 0.73 | 11 | 12 |
| BM MSC 9 | 0.14 | 0.46 | 0.93 | 0.92 | 10 | 7 |
| BM MSC 10 | 0.1 | 0.53 | 0.57 | 0.91 | 7 | 6 |
| BM MSC 11 | 0.09 | 0.79 | 0.65 | 0.95 | 7 | 5 |
| BM MSC 4 | 0.1 | 0.6 | 0.58 | 0.94 | 9 | 5 |
| BM MSC 12 | 0.06 | 0.7 | 0.32 | 0.93 | 3 | 4 |
| BM MSC 12 | 0.07 | 0.67 | 0.38 | 0.92 | 5 | 4 |
| BM MSC 12 | 0.08 | 0.68 | 0.37 | 0.91 | 7 | 5 |
| BM MSC 13 | 0.02 | 0.6 | 0.36 | 0.96 | 4 | 3 |
| BM MSC 13 | 0.03 | 0.53 | 0.51 | 0.81 | 8 | 10 |
| BM MSC 13 | 0.1 | 0.43 | 0.65 | 0.82 | 11 | 10 |
| BM MSC 14 | 0.08 | 0.64 | 0.4 | 0.9 | 4 | 5 |
| BM MSC 14 | 0.17 | 0.57 | 0.54 | 0.79 | 8 | 9 |
| BM MSC 14 | 0.2 | 0.48 | 0.61 | 0.72 | 10 | 12 |
| BM MSC 15 | 0.07 | 0.7 | 0.28 | 0.88 | 3 | 5 |
| BM MSC 15 | 0.09 | 0.67 | 0.4 | 0.9 | 5 | 5 |
| BM MSC 15 | 0.37 | 0.42 | 0.61 | 0.7 | 9 | 11 |
| BM MSC 16 | 0.16 | 0.75 | 0.28 | 0.91 | 0 | 3 |
| BM MSC 16 | 0.06 | 0.81 | 0.19 | 0.93 | 2 | 3 |
| BM MSC 17 | 0.19 | 0.64 | 0.65 | 0.88 | 5 | 6 |
| BM MSC 17 | 0.07 | 0.53 | 0.54 | 0.75 | 9 | 12 |
| BM MSC 18 | 0.08 | 0.82 | 0.23 | 0.92 | 0 | 3 |
| BM MSC 18 | 0.23 | 0.72 | 0.72 | 0.8 | 12 | 9 |
| BM MSC 19 | 0.14 | 0.64 | 0.71 | 0.59 | 16 | 18 |
| BM MSC 19 | 0.12 | 0.6 | 0.74 | 0.57 | 19 | 20 |
| HUVEC 5 | 0.05 | 0.61 | 0.32 | 0.83 | 3 | 8 |
| HUVEC 5 | 0.41 | 0.19 | 0.77 | 0.45 | 19 | 22 |
| HUVEC 6 | 0.05 | 0.61 | 0.26 | 0.86 | 4 | 6 |
| HUVEC 6 | 0.27 | 0.22 | 0.56 | 0.45 | 17 | 22 |
| HUVEC 7 | 0.06 | 0.62 | 0.36 | 0.88 | 4 | 6 |
| HUVEC 7 | 0.11 | 0.24 | 0.58 | 0.5 | 12 | 22 |
| BM MSC 4 | 0.11 | 0.42 | 0.46 | 0.96 | 11 | 3 |
| BM MSC 5 | 0.17 | 0.46 | 0.66 | 0.75 | 11 | 12 |
| BM MSC 5 | 0.10 | 0.31 | 0.67 | 0.86 | 12 | 8 |
| BM MSC 8 | 0.07 | 0.18 | 0.37 | 0.75 | 14 | 11 |

| | | | | | | |
|---|---|---|---|---|---|---|
| BM MSC 1 | 0.08 | 0.44 | 0.44 | 0.94 | 10 | 4 |
| BM MSC 1 | 0.09 | 0.46 | 0.35 | 0.89 | 11 | 5 |
| BM MSC 3 | 0.14 | 0.27 | 0.65 | 0.69 | 13 | 15 |
| BM MSC 3 | 0.15 | 0.25 | 0.71 | 0.66 | 14 | 16 |
| BM MSC 3 | 0.17 | 0.31 | 0.69 | 0.58 | 15 | 19 |
| BM MSC 3 | 0.18 | 0.33 | 0.66 | 0.64 | 16 | 16 |
| BM MSC 2 | 0.13 | 0.61 | 0.44 | 0.77 | 12 | 10 |
| BM MSC 2 | 0.12 | 0.51 | 0.40 | 0.79 | 13 | 9 |
| BM MSC 2 | 0.25 | 0.49 | 0.44 | 0.68 | 14 | 12 |
| AT MSC 1 | 0.18 | 0.53 | 0.58 | 0.68 | 11 | 14 |
| AT MSC 1 | 0.21 | 0.49 | 0.61 | 0.70 | 12 | 13 |
| AT MSC 1 | 0.17 | 0.44 | 0.64 | 0.66 | 13 | 15 |
| AT MSC 1 | 0.19 | 0.53 | 0.62 | 0.72 | 14 | 12 |
| AT MSC 2 | 0.22 | 0.53 | 0.61 | 0.73 | 14 | 12 |
| AT MSC 2 | 0.31 | 0.51 | 0.70 | 0.75 | 15 | 10 |
| AT MSC 2 | 0.38 | 0.44 | 0.66 | 0.68 | 16 | 12 |
| AT MSC 2 | 0.36 | 0.49 | 0.66 | 0.70 | 17 | 11 |
| AT MSC 2 | 0.56 | 0.48 | 0.63 | 0.62 | 19 | 12 |
| AT MSC 3 | 0.10 | 0.44 | 0.46 | 0.74 | 10 | 12 |
| AT MSC 3 | 0.12 | 0.35 | 0.50 | 0.75 | 11 | 11 |
| AT MSC 3 | 0.11 | 0.36 | 0.51 | 0.68 | 12 | 14 |
| AT MSC 3 | 0.15 | 0.38 | 0.56 | 0.71 | 13 | 13 |
| HUVEC 1 | 0.37 | 0.54 | 0.88 | 0.65 | 10 | 15 |
| HUVEC 1 | 0.44 | 0.59 | 0.83 | 0.64 | 11 | 14 |
| HUVEC 1 | 0.67 | 0.47 | 0.71 | 0.73 | 12 | 7 |
| HUVEC 1 | 0.52 | 0.46 | 0.62 | 0.63 | 13 | 12 |
| HUVEC 1 | 0.65 | 0.60 | 0.81 | 0.61 | 15 | 13 |
| HUVEC 2 | 0.48 | 0.60 | 0.73 | 0.69 | 10 | 11 |
| HUVEC 2 | 0.65 | 0.56 | 0.67 | 0.53 | 11 | 15 |
| HUVEC 3 | 0.56 | 0.67 | 0.93 | 0.38 | 18 | 24 |
| HUVEC 3 | 0.54 | 0.64 | 0.94 | 0.36 | 19 | 24 |
| HUVEC 3 | 0.54 | 0.60 | 0.90 | 0.34 | 20 | 25 |
| HUVEC 3 | 0.65 | 0.53 | 0.93 | 0.28 | 22 | 26 |
| HUVEC 4 | 0.67 | 0.76 | 0.92 | 0.37 | 18 | 22 |
| HUVEC 4 | 0.62 | 0.71 | 0.93 | 0.28 | 19 | 27 |
| HUVEC 4 | 0.56 | 0.70 | 0.94 | 0.27 | 20 | 28 |
| HUVEC 4 | 0.52 | 0.76 | 0.91 | 0.27 | 22 | 28 |

Methylation values of single CpGs are depicted as β-values ranging from 0 to 1.

**Supplemental Table S7. BBAseq primers**

| gene name | primer | Sequence 5’ -> 3’ |
|---|---|---|
| *ALOX12* | forward | CTCTTTCCCTACACGACGCTCTTCCGATCTGGGGTTATTTTTAATTTTTAAAGGAT |
| *ALOX12* | reverse | CTGGAGTTCAGACGTGTGCTCTTCCGATCTAAAACACAACCAATCCCCACAA |
| *DOK6* | forward | CTCTTTCCCTACACGACGCTCTTCCGATCTATGAGAAAATTGAGATATAATTTTATTTAGGAAATAG |
| *DOK6* | reverse | CTGGAGTTCAGACGTGTGCTCTTCCGATCTCACCTTTTCTTCTAAAATCTACAAATCCC |
| *LTC4S* | forward | CTCTTTCCCTACACGACGCTCTTCCGATCTTTTTAGGGTTTTGTAGATTTTTATATTATGTTGGAGTTAG |
| *LTC4S* | reverse | CTGGAGTTCAGACGTGTGCTCTTCCGATCTCACCCAAAAAACCTTAAACAAATTTCC |
| *TNNI3K* | forward | CTCTTTCCCTACACGACGCTCTTCCGATCTATTTTGTGGTTTTTTATAATGTTTTAGGAGTGTGATAA |
| *TNNI3K* | reverse | CTGGAGTTCAGACGTGTGCTCTTCCGATCTCCATAATCACTTTATTCACTACATCACCAATACCCATTC |
| *GRM7* | forward | CTCTTTCCCTACACGACGCTCTTCCGATCTTTGGGATTATTGTTGATTT |
| *GRM7* | reverse | CTGGAGTTCAGACGTGTGCTCTTCCGATCTCCCCTACTACCTACTAAAAATA |
| *CASR* | forward | CTCTTTCCCTACACGACGCTCTTCCGATCTTGTAATAGGTATTTGGTTGTAGT |
| *CASR* | reverse | CTGGAGTTCAGACGTGTGCTCTTCCGATCTCCCAAACTCTTACTCATTCTA |
| *PRAMEF2* | forward | CTCTTTCCCTACACGACGCTCTTCCGATCTTTTGAGGGTATTTAGAAGAGAT |
| *PRAMEF2* | reverse | CTGGAGTTCAGACGTGTGCTCTTCCGATCTTCCCTAACTAACTAACTACTAATC |
| *SELP* | forward | CTCTTTCCCTACACGACGCTCTTCCGATCTAGAAGGTAGAAAATTAGTAGAGTT |
| *SELP* | reverse | CTGGAGTTCAGACGTGTGCTCTTCCGATCTCAACATAAAACTCCATAACTA |
| *CASP14* | forward | CTCTTTCCCTACACGACGCTCTTCCGATCTTTGGAGATTTAGTGAGATAATA |
| *CASP14* | reverse | CTGGAGTTCAGACGTGTGCTCTTCCGATCTAACAAAACAAATAACCCATATA |
| *KRTAP13.3* | forward | CTCTTTCCCTACACGACGCTCTTCCGATCTGAGATTTGTTGGAGGTTTAA |
| *KRTAP13.3* | reverse | CTGGAGTTCAGACGTGTGCTCTTCCGATCTCCCAATAAAAAACAACTCC |

**Supplemental Table S8. Hairpin linkers**

| Name | Sequence 5' -> 3' | CpG sites [enzyme] |
|---|---|---|
| Hairpin-1 | Phosphate-CTAGCGATGCDDDDDDDDGCATCGCT | *CASR* [AccI] |
| Hairpin-2 | Phosphate-TAAGCGATGCDDDDDDDDGCATCGCT | *SELP, PRAMEF2, GRM7, GNAS, DOK6* [CviQI] |
| Hairpin-3 | Phosphate-TGAAGCGATGCDDDDDDDDGCATCGCT | *KRTAP13.3* [DdeI] |
| Hairpin-4 | Phosphate-AGAGCGATGCDDDDDDDDGCATCGCT | *C12orf12, TNNI3K* [AccI] |
| Hairpin-5 | Phosphate-TTAAGCGATGCDDDDDDDDGCATCGCT | *LTC4S* [DdeI] |

Hairpin linkers were designed with different overhangs, which are complementary to the sticky ends of the restriction digest at each CpG site with respective enzymes indicated in square brackets. Unique molecular identifiers (UMIs) consisted of random A,G and T nucleotides represented by the letter D.

**Supplemental Table S9. Hairpin primers**

| Primer name | Sequence 5' -> 3' |
|---|---|
| HPnewCASR forw | GTTTAAATTTTTATTTATTTTGTAAGATTTAGG |
| HPnewCASR rev | ACTCTTACTCATTCTACAAAACTC |
| HPnewGRM7 forw | GAGTAGTATGGTTTAGTTGAGG |
| HPnewGRM7 rev | CATAATCCAACTAAAAAAACTACTCC |
| HPnewKRTAP13.3 forw | GTAATTTTTGTTTGATTATGTATGTTGG |
| HPnewKRTAP13.3 rev | ATATTAAATCCAACCCCTACCAC |
| HPnewPRAMEF2 forw | GGTTGGTTGTTGATTAGATGGG |
| HPnewPRAMEF2 rev | CTAACTACTAATCAAATAAACATAACCC |
| HPnewSELP forw | TAGGTAAAGGTTTAGAAAGTGAGG |
| HPnewSELP rev | TAAACAAAAAACAAAAAACCAACAAAATCAC |
| HPnewDOK6 forw | TTAAAGAGATATAATAAAAATAGGTGGG |
| HPnewDOK6 rev | CAAAAAACTACTTAAAATACCTATTTCAC |
| HPnewLTC4S forw | GTTTTTTGTTTTTATTTAGGTTGTTTTTGG |
| HPnewLTC4S rev | ATCCAAACTATTCCTAACAACCC |
| HPnewTNNI3K forw | GTATTATTAGTATTTATTTTTAGTAGAGTG |
| HPnewTNNI3K rev | ATTCACTACATCACCAATACCC |
| HPnewGNAS forw | ATTTTTTTTTTTTGTTTAGAGAGG |
| HPnewGNAS rev | CATCCCTTCTTCTTACTC |
| HPnewC12orf12 forw | AGTTTTAGTTTTATTTAGTATATTTGGG |
| HPnewC12orf12 rev | AATCCCACCCAACACACCT |

**Supplemental Table S10. 4C primers**

| Viewpoint | primer | sequence |
|---|---|---|
| *ALOX12* | forward | ATCCAGTAAGGGACAGACAC |
| *ALOX12* | reverse | GTAATATCCAAATAAAATGGCTC |
| *LTC4S* | forward | GGGTCCTCCCATGGAGAATT |
| *LTC4S* | reverse | GAGCACCATGAAGAACTTTGC |
| *KRTAP13.3* | forward | CACACACCATTGAAATGACAG |
| *KRTAP13.3* | reverse | CTCTGCAATTCTTGCCTGAC |
| *CASR* | forward | AGACCGTGACCTTGGCATAG |
| *CASR* | reverse | ATGCAGTATTCCACCCTTGC |
